# Cyclic-di-GMP is required for corneal infection by *Pseudomonas aeruginosa* and modulates host immunity

**DOI:** 10.1101/098749

**Authors:** Joey Kuok Hoong Yam, Thet Tun Aung, Song Lin Chua, Yingying Cheng, Gurjeet Singh Kohli, Jianuan Zhou, Florentin Constancias, Yang Liu, Zhao Cai, May Margarette Santillan Salido, Daniela I. Drautz-Moses, Scott A. Rice, Stephan Christoph Schuster, Bin Wu, Staffan Kjelleberg, Tim Tolker-Nielsen, Roger W. Beuerman, Michael Givskov, Liang Yang

## Abstract

Biofilms are extremely tolerant toward antimicrobial treatment and host immune clearance due to their distinct physiology and protection by extracellular polymeric substances. Bis-(3´-5´)-cyclic dimeric guanosine monophosphate (c-di-GMP) is an essential messenger that regulates biofilm formation by a wide range of bacteria. However, there is a lack of physiological characterization of biofilms *in vivo* as well as the roles of c-di-GMP signaling in mediating host-biofilm interactions. Here, we employed dual RNA-Seq to characterize the host and pathogen transcriptomes during *Pseudomonas aeruginosa* infection using a mouse keratitis model. *In vivo P. aeruginosa* biofilms maintained a distinct physiology compared with *in vitro P. aeruginosa* biofilms, with enhanced virulence and iron uptake capacity. C-di-GMP synthesis was enhanced in *P. aeruginosa* cells *in vivo,* potentially due to down-regulation of the expression of several phosphodiesterases (e.g., DipA, NbdA). Increased intracellular c-di-GMP levels were required for long-term ocular colonization of *P. aeruginosa* and impaired host innate immunity.

## Introduction

Bacterial pathogens are able to form surface-attached, complex-multicellular biofilm communities in clinical settings, including hospital environments and the site of host infection [1]. Bacterial cells within biofilms are encased in extracellular polymeric substances (EPS), which mainly consist of extracellular proteins, exopolysaccharides and extracellular DNA (eDNA) [2-4]. These densely packed EPS serve as a protective shield that reduces the penetration of antimicrobials as well as phagocytes, thus greatly enhancing the survival of pathogens under hostile environmental conditions [3,5]. In addition to EPS, the specific physiology and social behaviors of biofilm cells also contribute to their tolerance toward antimicrobials and host immune clearance [6-8].

Biofilm formation has been proposed to be the main cause of a wide range of persistent infections, including well-characterized cystic fibrosis lung infections [9], dental plaque-induced gingival diseases [10] and urinary tract infections [11]. However, there is a lack of physiological characterization for the presence and/or impact of biofilms in a number of infections. One example is ocular infection, which continues to be the main cause of ocular disease and vision loss [12]. Despite increasing evidence that bacterial biofilms play a role in ocular infections, most studies have been based on *in vitro* characterizations of the biofilm formation capacity of ocular bacterial isolates [13]. Studies of bacterial physiology and host-pathogen interactions within the ocular tissue might provide mechanistic insights into the pathogenesis of bacterial ocular infections.

The Gram-negative opportunistic pathogen *Pseudomonas aeruginosa* is an important etiologic agent of a variety of ocular infectious diseases [14]. Current evidence indicates that the induction of the T helper 1 (Th1)-type immune response in C57BL/6 mice does not eradicate *P. aeruginosa* during corneal infections, and contributes to corneal destruction and perforation associated with the mouse keratitis model [15]. The inability of the corneal innate immune response to clear *P. aeruginosa* infections leads to the hypothesis that biofilm formation protects *P. aeruginosa* cells against immune clearance during corneal infections. As a model organism for biofilm research, the biofilm formation mechanisms of *P. aeruginosa* have been well characterized [16]. Biofilm formation and the dispersal of *P. aeruginosa* is regulated by the intracellular secondary messenger c-di-GMP, which is synthesized by diguanylate cyclase (DGC) enzymes and degraded by phosphodiesterase (PDE) enzymes [17]. A high intracellular content of c-di-GMP represses *P. aeruginosa* cell motility and induces polysaccharide (Pel, Psl and alginate) synthesis, resulting in biofilm formation *in vitro* [18,19]. Lowering the intracellular c-di-GMP content of biofilm cells by inducing the expression of PDE enzymes have been shown to disperse biofilms both *in vitro* [20] and *in vivo* [21]. However, it is unclear whether c-di-GMP regulates *P. aeruginosa* biofilm formation and host-pathogen interactions during ocular infections.

In the present study, we employed dual RNA-Seq technology for comparative analysis of the transcriptomes of *in vivo P. aeruginosa* with different c-di-GMP contents, and we provide evidence that c-di-GMP synthesis is induced *in vivo* and contributes to biofilm formation and host-pathogen interactions during mouse corneal infection.

## Materials and Methods

### Ethics statement

All animal experiments were conducted in compliance with the ARVO statement for the Use of Animals in Ophthalmic and Vision Research, the Guide for the Care and Use of Laboratory Animals (National Research Council) under SingHealth Institutional Animal Care and Use Committee (IACUC) protocol number #2014/SHS/901, SingHealth Institutional Biosafety Committee (IBC) approval number SHSIBC-2014-015 and under the supervision of SingHealth Experimental Medical Centre (SEMC).

### Bacterial strains and culture media

The bacterial strains (Table 1) were routinely cultivated in Luria-Bertani (LB) medium. For marker selection in *P. aeruginosa*, 30 μg ml^−1^ gentamicin (Gm), 50 μg ml^−1^ tetracycline (Tc), 100 μg ml^−1^ streptomycin (Strep) or 200 μg ml^−1^ carbenicillin (Cb) were used, as appropriate.

**Table 1.**
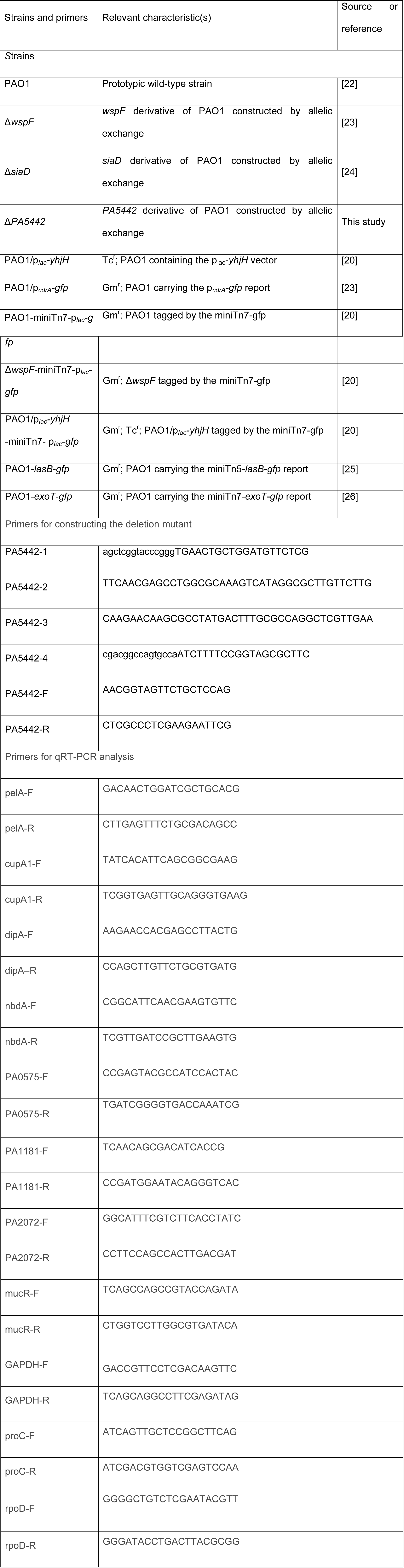
Strains and primers used in this study.

### Mouse infection experiment

Female C57BL6 (6-8 weeks old, 20-30 g) mice (The Jackson Laboratory) were anesthetized subcutaneously with 100 mg kg^−1^ ketamine and 10 mg kg^−1^ xylene and placed under a stereoscopic microscope. A sterile miniblade (Beaver-Visitec International, MA, USA) was used to make corneal scratches (n=3, each 1 mm long) that did not breach the superficial stroma on the right eye, while the left eye remained untouched [27,28]. Ten microliters of a bacterial suspension containing 1 × 10^5^ colony forming units (CFU)/μl *P. aeruginosa* was topically applied to infect the scratched cornea.

The mouse corneas were infected with the PAO1-*lasB-gfp* and PAO1-*exoT-gfp* strains for 2 days. The corneas were then dissected and stained with 20 μl of 5 μM SYTO®62 Red Fluorescent Nucleic Acid Stain (Thermo Fisher Scientific, Singapore), which stains nucleic acids in both bacterial and corneal cells, for 15 min. They were then visualized under a Zeiss LSM780 Confocal Laser Scanning Microscope, CLSM (Carl Zeiss, Jena, Germany) with a 63x objective lens using an argon laser (488 nm, 535 nm emission) for GFP observation and a helium laser (633 nm, 680 nm) for STYO®62. For the *in vitro* biofilm controls, the PAO1-*lasB-gfp* and PAO1-*exoT-gfp* strains were grown at 37°C in ABT minimal medium [29] supplemented with 2 gl^−1^ glucose and 2 gl^−1^ casamino acids (ABTGC) using the μ-Slide 8-well imaging chamber (Ibidi GmbH, Martinsried, Germany) for 18 hours under static conditions. Similarly, the *in vitro* biofilms were stained with SYTO®62 and imaged under a CLSM.

The PAO1/P_*cdrA*_-*gfp* and PAO1/p_*lac*_-*gfp* cells were used for the corneal infections to observe the expression of GFP. Infected corneas were dissected at 2 hours post-infection (hpi), 4 hpi and 8 hpi as well as at 1 day post-infection (dpi), 2 dpi, 4 dpi and 7 dpi. As mentioned above, the dissected corneas were stained with SYTO®62 and viewed under a CLSM to evaluate the presence of GFP expression and the total biomass. Planktonic PAO1/P_*cdrA*_-*gfp* and PAO1/p_*lac*_-*gfp* were used as controls. The bacterial cells were cultivated in LB medium for 6 h (mid-log phase) with shaking at 37°C and observed under a CLSM.

### Construction of the *P. aeruginosa PA5442* mutant

The upstream and downstream sequences of *PA5442* were obtained from the *Pseudomonas* genome database (http://www.pseudomonas.com/), and the primers used for amplification of the *PA5442* are listed in Table 1. Briefly, the up and down flanking fragments of *PA5442* were amplified with primer pairs PA5442-1 & PA5442-2, and PA5442-3 & PA5442-4, respectively. The PCR products were purified and assembled with *Bam*HI and *Hind*III-digested pK18 (Gm^r^) plasmid using Gibson Assembly Master Mix (NEB), and then transformed into *E. coli* DH5α competent cells. Positive transformants were picked and grown in LB medium supplemented with 30 μg/ml Gm. In-frame deletion mutant was generated by triparental conjugation with the aid of RK600 [30]. Mutants were confirmed by amplification using primers PA5442-F and PA5442-R and sequencing. All other mutants were generated in the same manner.

### Colonizing pattern and immune response characterization

Female C57BL6 mice (6-8 weeks old, 20-30 g) were infected with the p*_lac_*-*gfp* tagged *P. aeruginosa* PAO1 wild-type, Δ*wspF* mutant or the PAO1/p*_lac_*-*yhjH* strain as described above. To visualize the presence of bacteria and characterize the colonization patterns and recruitment of phagocytes, the infected corneas were dissected at 2 dpi and 7 dpi and stained for 15 min with 20 μl of solution consisting of 1 μM of LysoTracker® Red DND-99 and approximately 0.165 μM Alexa Fluor® 635 phalloidin (Life Technologies) (633 nm, emission 647), which stain phagocytes and F-actin of eukaryotic cells, respectively, prior to CLSM imaging.

To measure the cytokines, infected corneas at 2 dpi and 7 dpi were dissected and homogenized using a VCX 750 probe sonicator (Sonics & Materials, US). The cell debris was removed by centrifugation at 13,000 × *g* for 5 min, and the supernatants were used to characterize the innate immune response. The cytokine levels were normalized to the total proteins using a Qubit® 2.0 Fluorometer (Invitrogen) prior to characterizing the pro-inflammatory response using the Bio-Plex Pro™ Mouse Cytokine 8-plex Assay (Bio-Rad) with the Bio-Plex® 200 system (Bio-Rad).

### Transcriptomic analysis

#### RNA preparation

Female C57BL6 mice (6-8 weeks old, 20-30 g) were infected with *P. aeruginosa* PAO1 wild-type as described above. Infected corneas were dissected at 2 dpi and 7 dpi, and total RNA was extracted using the MiRNeasy mini kit (Qiagen). A vigorous Turbo DNA-free protocol was used for DNase treatment (Ambion). The integrity of the total RNA and the level of DNA contamination were assessed with an Agilent 2200 Tapestation (Agilent Technologies) and a Qubit 2.0 Fluorometer (Invitrogen). Three biological replicates were used for the transcriptomic analysis at 2 dpi and 7 dpi, respectively.

#### RNA sequencing and data analysis

Gene expression analysis was conducted by Illumina RNA sequencing (RNA-Seq technology). RNA-Seq was conducted for three biological replicates of each sample. Libraries were produced using an Illumina TruSeq Stranded messenger RNA Sample Prep Kit. The libraries were sequenced using the Illumina HiSeq 2500 platform with a paired-end protocol and read lengths of 100 nt.

A total of approximately 20 million reads were obtained for each sample. Raw reads were trimmed, adaptor sequences removed and any reads below 50 bp discarded. Trimmed reads were then mapped onto the mouse reference genome, which can be downloaded from the Ensemble database (ftp.ensembl.org) using the ‘RNA-Seq and expression analysis’ application of the CLC genomics Workbench 9.0 (CLC Bio, Aarhus, Denmark). The following criteria were used to filter the unique sequence reads: maximum number of hits for a read of 1, minimum length fraction of 0.9, minimum similarity fraction of 0.8 and maximum number of mismatches of 2. A constant of 1 was added to the raw transcript count value to avoid any problems caused by 0.

The raw count table of transcripts was then used as an input for the Deseq2 R package for differential expression analysis [31]. First, the raw counts were normalized according to the sample library size. Next, a negative binomial test was performed to identify the differentially expressed genes. The transcripts were determined as differentially expressed among pairwise comparisons when their absolute fold-change value was greater than five and the associated adjusted P-value was smaller than 0.05. The normalized transcripts were then log2(N+1) transformed prior to principal component analysis and UPGMA hierarchical clustering for the sample dendrogram on the heatmap.

### qRT–PCR analysis

Total RNA was extracted using a miRNeasy Mini Kit (Cat. No. 74104, Qiagen) with on-column DNase digestion. The purity and concentration of the RNA were determined by NanoDrop spectrophotometry, and the integrity of the RNA was measured using an Agilent 2200 TapeStation System. The contaminating DNA was eliminated using a Turbo DNA-free Kit (Cat. No. AM1907, Ambion) and confirmed by real-time PCR amplification of the *rpoD* gene using total RNA as the template.

Quantitative reverse transcriptase PCR (qRT-PCR) was performed using a two-step method. First-strand cDNA was synthesized from total RNA using the SuperScript III First-Strand Synthesis SuperMix kit (Cat. No. 18080-400, Invitrogen). The cDNA was used as a template for qRT–PCR with a SYBR Select Master Mix kit (Cat. No. 4472953, Applied Biosystems by Life Technologies) on an Applied Biosystems StepOnePlus Real-Time PCR System. The *rpoD*, *proC* and GAPDH genes were used as endogenous controls. All pairs of primers were confirmed to have an efficiency between 90-110% before performing the qRT-PCR. Melting curve analyses were employed to verify the specific single-product amplification.

### Accession codes

The dual-species RNA-Seq data have been deposited in the NCBI Short Read Archive (SRA) database under accession code SRP078905. The RNA-Seq data for *P. aeruginosa in vitro* planktonic cells, biofilm cells and dispersed cells have been previously published [29] and deposited in the NCBI Short Read Archive (SRA) database under accession code SRP041868.

## Results

### Transcriptomes of in vivo P. aeruginosa biofilms

To examine the physiology of *in vivo P. aeruginosa* biofilms and their impact on the host, we performed dual RNA-Seq analysis of biofilms formed by *P. aeruginosa* PAO1 wild-type, its high intracellular c-di-GMP containing derivative Δ*wspF* mutant and its intracellular c-di-GMP-depleted derivative PAO1/p_*lac*_-*yhjH* mutant at 2 day-post-infection (dpi) and 7 dpi in the mouse model of experimental keratitis infection, which has recently been suggested to be a biofilm-associated infection according to imaging-based characterization [32]. The Δ*wspF* mutation, which encodes a repressor for WspG DGC, is frequently observed in *P. aeruginosa* clinical isolates [33]. We used the heterologous PDE YhjH to deplete the *P. aeruginosa* intracellular c-di-GMP content [34], which presumably is not subject to regulation in *P. aeruginosa* since it contains only an EAL domain and is from *E. coli.*

Both *P. aeruginosa* and mouse corneal RNA were purified from infected corneal samples, and RNA-Seq was performed using the Illumina HiSeq 2500 System. Between 404,204 and 1,192,838 reads were mapped to the *P. aeruginosa* genome from 2 dpi samples, while between 3,238 and 107,380 reads were mapped to the *P. aeruginosa* genome from 7 dpi samples (Table S1). Due to an insufficient number of reads, the *P. aeruginosa* samples from 7 dpi infected corneas were discarded. The transcriptomes of *P. aeruginosa* PAO1 wild type, Δ*wspF* mutant and PAO1/p_lac_-*yhjH* strain from 2 dpi infected corneas were compared to our previously characterized transcriptomes of *in vitro P. aeruginosa* PAO1 planktonic cells, biofilm cells and dispersed cells [29]. Projection of the transcriptomes by principal component analysis (PCA) revealed that *in vivo P. aeruginosa* cells have a distinct physiology compared with *P. aeruginosa* cells *in vitro*, which was largely independent of the intracellular c-di-GMP content (Figure 1A). The PCA analysis suggested that the *in vivo P. aeruginosa* cells has significant similarities to *P. aeruginosa* biofilm cells cultured *in vitro* (reflected by PC2). Using a negative binomial test with a *P*-value cut-off of 0.05 and a fold-change cut-off of 5, we found that 348 genes (Supplementary Data 1) were up-regulated and 301 genes (Supplementary Data 2) were down-regulated in *P. aeruginosa* cells cultured *in vivo* compared with *in vitro P. aeruginosa* biofilms. Genes responsible for iron uptake (such as *PA4836, phuR, hasAp*, and *hasR)* and type III secretion mechanisms (such as *exoS, pcrV, pscF*, and *exoT)* were highly expressed in *P. aeruginosa* cells *in vivo* compared with *P. aeruginosa* biofilm cells *in vitro* (Supplementary Data 1). In contrast, *in vitro P. aeruginosa* biofilm cells exhibited a higher expression level of genes involved in quorum sensing (e.g., *lasA, chiC, rhlB, lecB*, and *hcnB)* compared with *in vivo* biofilm *P. aeruginosa* cells (Supplementary Data 2). To confirm the RNA-Seq analysis, we used GFP-based reporter fusions of the *lasB* gene (quorum-sensing controlled) and *exoT* gene (type III secretion controlled) to measure the expression of quorum-sensing and type III secretion in *P. aeruginosa* cells *in vivo* and *P. aeruginosa* biofilm cells *in vitro.* As shown in Figure 1B, these results were consistent with the RNA-Seq analysis (Supplementary Data 1 and 2).

**Figure 1.**
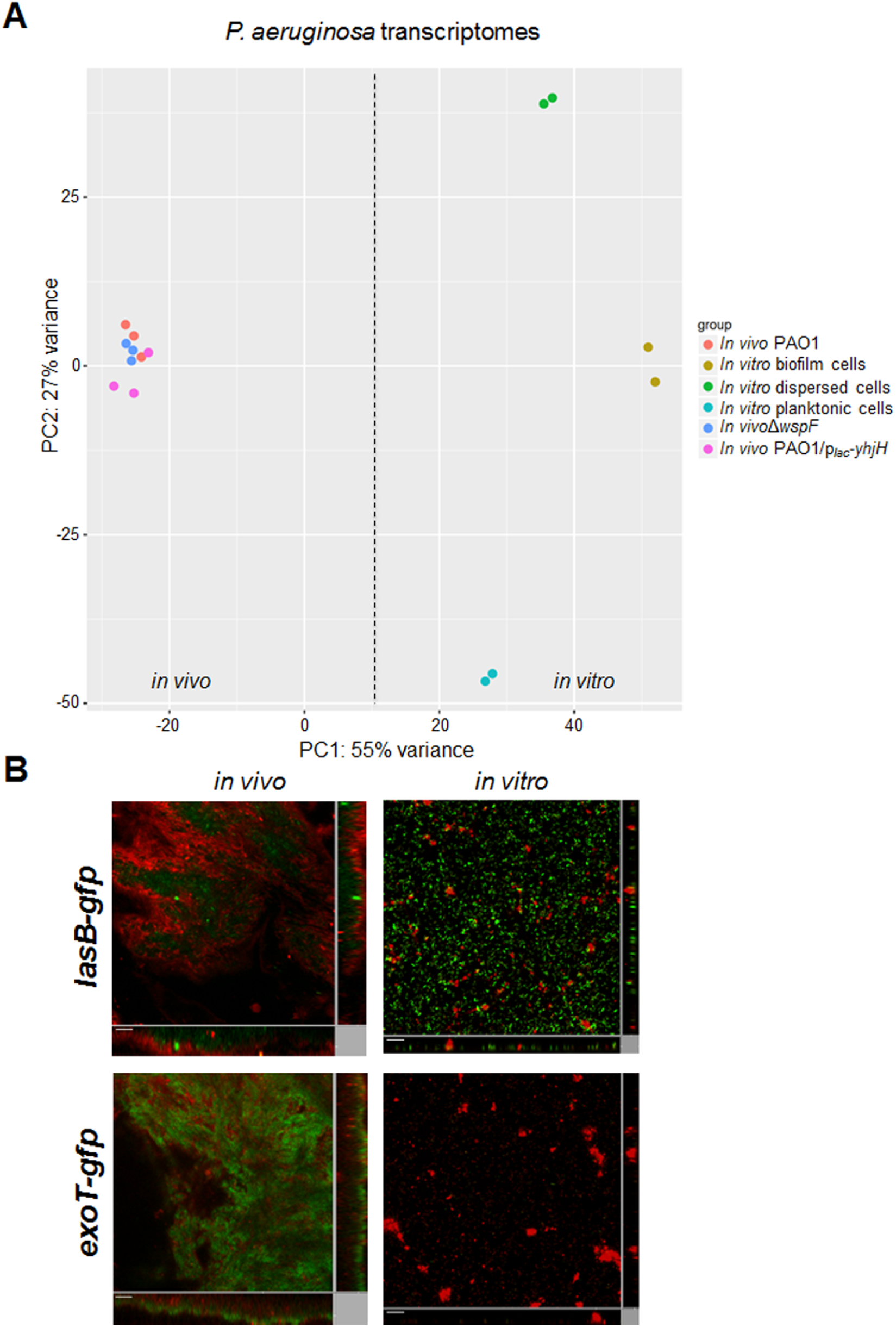
**A.** PCA of the transcriptomes of *in vitro P. aeruginosa* PAO1 planktonic cells, SNP-dispersed cells, biofilm cells and *in vivo P. aeruginosa* cells at 2 dpi of mouse corneas by PAO1, Δ*wspF* and PAO1/p_*lac*_-*yhjH*. Raw RNA-Seq data were normalized using the DESeq package before PCA analysis. The dotted line was used to separate *in vivo P. aeruginosa* cells and *in vitro P. aeruginosa* cells by PCA analysis. **B.** Confocal image of *in vitro P. aeruginosa* PAO1 biofilm cells and *in vivo P. aeruginosa* PAO1 cells at 2 dpi of mouse corneas. PAO1 strains are tagged with *lasB-gfp* (quorum-sensing reporter) and *exoT-gfp* (type III secretion reporter), respectively. The expression of reporter fusions was recorded by the green fluorescence, while the total biomass was imaged based on the SYTO®62 stain (red fluorescence). Representative images are shown for each condition. Scale bars, 10 μm.

### C-di-GMP synthesis is induced in P. aeruginosa during corneal infections

To further capture the “biofilm features” of the *in vivo P. aeruginosa* cells, we compared the transcriptomes of *in vivo P. aeruginosa* PAO1 cells with the *in vitro P. aeruginosa* PAO1 planktonic cells. Using a negative binomial test with a *P*-value cut-off of 0.05 and a fold-change cut-off of 5, we found that 330 genes (Supplementary Data 3) were up-regulated and 414 genes (Supplementary Data 4) were down-regulated in *in vivo P. aeruginosa* cells compared with *in vitro P. aeruginosa* planktonic cells. Genes encoding the biofilm matrix, such as Pel exopolysaccharide (e.g., *pelB, pelC, pelD*, and *pelF)* and cup fimbriae (e.g., *cupA1, cupA2)* were highly expressed in *in vivo* PAO1 cells compared with *in vitro* PAO1 planktonic cells (Supplementary Data 3), which was confirmed by qRT-PCR analysis (Figure S1).

Since the expression of genes involved in Pel exopolysaccharide and cup fimbriae synthesis is induced by a high intracellular c-di-GMP content in *P. aeruginosa* [35,36], we next examined whether the intracellular c-di-GMP content in *P. aeruginosa* increased during corneal infections by using a *P. aeruginosa* PAO1 strain carrying the p_*cdrA*_-*gfp* c-di-GMP reporter fusion to establish infections [23]. The *cdrA* gene encodes a large extracellular matrix adhesin, which is up-regulated in the presence of high levels of intracellular c-di-GMP [19]. Planktonic PAO1/p_*cdrA*_-*gfp* cells showed an extremely low expression level of the p_*cdrA*_-*gfp* fusion (below the detection threshold) compared with planktonic PAO1 control cells carrying p_*lac*_-*gfp* (Figure 2A and 2B), implying that the level of intracellular c-di-GMP content was low. This result correlates well with the previous study showing that expression of the p_*cdrA*_-*gfp* fusion could only be detected in planktonic *P. aeruginosa* cells with high intracellular c-di-GMP contents, such as the Δ*wspF* mutant [20,23]. A p*_lac_*-*gfp* tagged PAO1 strain showed a clear biofilm-like aggregation morphology at 8 hours post-infection (8 hpi) (Figure 2C). To capture the transition stage of p_*cdrA*_-*gfp* expression, we monitored the expression of the p_*cdrA*_-*gfp* fusion in the PAO1/p_*cdrA*_-*gfp* strain at 2 hpi, 4 hpi and 8 hpi, and found that gfp expression was induced in approximately half of the PAO1/p_*cdrA*_-*gfp* cells at 2 hpi (Figure 2D), suggesting that c-di-GMP synthesis in *P. aeruginosa* might be induced shortly after the establishment of corneal infections. At 4 hpi, only *gfp*-expressing cells were observed in the PAO1/p_*cdrA*_-*gfp*-infected corneas (Figure 2E), suggesting that bacteria with low intracellular c-di-GMP contents might be eradicated by the host immune system due to their low fitness. At 8 hpi, small biofilm-like aggregates were visualized in the PAO1/p_*cdrA*_-*gfp*-infected corneas with bright GFP fluorescent signals (Figure 2F). Expression of the p_*cdrA*_-*gfP* fusion was maintained at a high level in mouse corneal samples at 2 dpi and 4 dpi, suggesting a constitutive increase in c-di-GMP content in *P. aeruginosa* cells during the course of corneal infection (Figure 2G and 2H). Interestingly, expression of the p_*cdrA*_-*gfp* fusion was reduced in PAO1 cells at 7 dpi (Figure 2I). No background *gfp* fluorescence could be detected in the control scratched mouse corneas (Figure S2).

**Figure 2.**
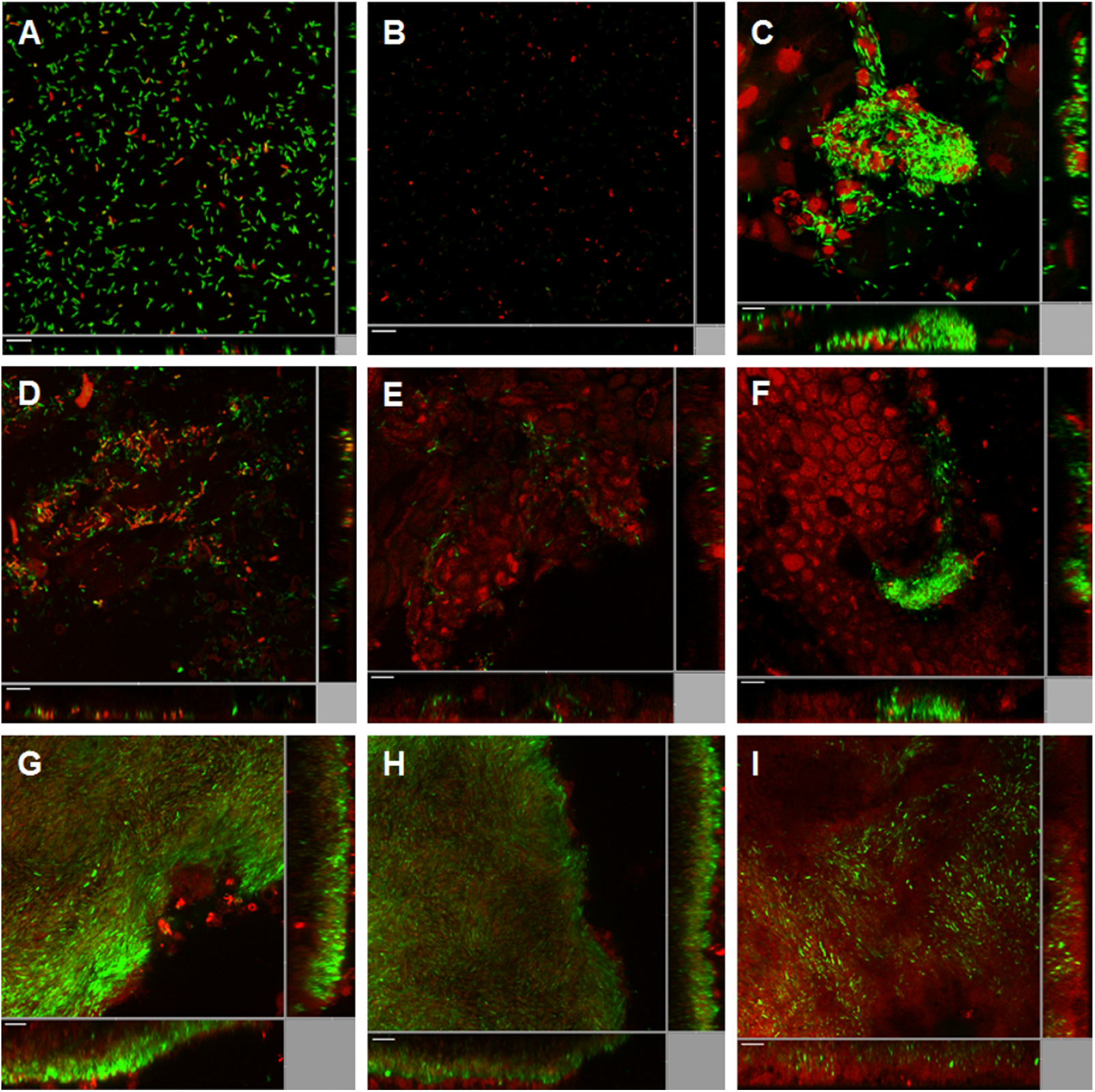
Induction of the c-di-GMP reporter fusion p_*cdrA*_-*gfp* in *P. aeruginosa* PAO1 during mouse corneal infection. **A.** Planktonic PAO1/p_*lac*_-*gfp* cells. **B.** Planktonic PAO1/p_*cdrA*_-*gfp* cells. **C.** PAO1/p_*lac*_-*gfp* cells at 8 hours post-infection (8 hpi) of mouse cornea. **D.** PAO1/p_*cdrA*_-*gfp* cells at 2 hpi of mouse cornea. **E**. PAO1/p_*cdrA*_-*gfp* cells at 4 hpi of mouse cornea. **F.** PAO1/p_*cdrA*_-*gfp* cells at 8 hpi of mouse cornea. **G.** PAO1/p_*cdrA*_-*gfp* cells at 1 day post-infection (1 dpi) of mouse cornea. **H.** PAO1/p_*cdrA*_-*gfp* cells at 4 dpi of mouse cornea. **I.** PAO1/p_*cdrA*_-*gfp* cells at 7 dpi of mouse cornea. SYTO®62 was used to stain host cells as well as *P. aeruginosa* cells lacking fluorescence. Green fluorescence represents constitutive expression of p_*lac*_-*gfp* (A, C) and expression of the p_*cdrA*_-*gfp* reporter fusion (B, D - I), and red fluorescence represents SYTO®62 staining. Representative images are shown for each condition. Scale bars, 10 μm.

To identify the potential DGCs responsible for the increased intracellular c-di-GMP content of *P. aeruginosa* during ocular infections, we first examined the expression of p_*cdrA*_-*gfP* in the *PA0169 (siaD)* and *PA5442* (an uncharacterized gene encoding a GGDEF domain-containing protein) mutants during ocular infection because these two DGC (or potential DGC)-encoding genes were shown to be induced in *in vivo P. aeruginosa* cells from ocular infection by 2.8 and 2.6-fold, respectively, compared with planktonic cells (Supplementary Data 3). However, the p_*cdrA*_-*gfp* fusion was expressed during corneal infection in both mutants (Figure S3), suggesting that the induction of c-di-GMP reporter p_*cdrA*_-*gfp* expression was unlikely to be due to the activity of these DGCs.

By examining the transcriptomic data of *in vivo P. aeruginosa* cells in comparison with planktonic cells, we noticed that the expression of some previously characterized PDE-encoding genes, *dipA (PA5017), mucR (PA1727), nbdA (PA3311), PA0575, PA1181*, and *PA2072*, was down-regulated in *in vivo P. aeruginosa* cells compared with planktonic cells (Supplementary Data S4), which was confirmed by qRT-PCR analysis (Figure S1). The down-regulation of these PDE-encoding genes might be responsible for the increased intracellular c-di-GMP content of *in vivo P. aeruginosa* cells, which again suggests that corneal infection caused by *P. aeruginosa* is a biofilm-associated infection.

### C-di-GMP is required for the establishment of microcolonies of P. aeruginosa during corneal infection

Neutrophil recruitment following *P. aeruginosa* corneal infection has been well documented [37]. However, the recruited neutrophils are unable to eradicate the *P. aeruginosa* during corneal infection, likely due to the high level of tolerance of *P. aeruginosa* biofilms toward host phagocytic cells. Both *in vitro* and *in vivo* experiments provide evidence that host phagocytic cells are unable to penetrate *P. aeruginosa* biofilms [38,39]. Since the comparative transcriptomic analysis and c-di-GMP reporter fusion p_*cdrA*_-*gfp* assay both suggest that c-di-GMP synthesis is induced during *P. aeruginosa* ocular infections, we hypothesized that increased c-di-GMP content is required for *in vivo P. aeruginosa* colonization.

We monitored the colonization patterns of p_*lac*_-*gfp*-tagged *P. aeruginosa* PAO1, the Δ*wspF* mutant (with high c-di-GMP content) and the PAO1/p_*lac*_-*yhjH* mutant (with c-di-GMP depleted) at 2 dpi and 7 dpi. To visualize the interaction between the host immune cells and *P. aeruginosa* biofilms, the LysoTracker® Red DND-99 stain (Molecular Probes, USA) was used to identify immune cells in C57BL6 mouse corneas infected with *P. aeruginosa.* LysoTracker® Red DND-99 stains the acidic lysosomes of host immune cells such as macrophages and neutrophils. Alexa Fluor® 635 phalloidin was also used to stain the F-actin of eukaryotic cells. As expected, *P. aeruginosa* PAO1 and the Δ*wspF* mutant formed large and densely packed microcolonies at the sites of infection, and they were surrounded by phagocytes at 2 dpi (Figure 3A, B) and 7 dpi (Figure 3D, E). This observation is very similar to the sites of *P. aeruginosa* infection in the lungs of cystic fibrosis patients [40], where *P. aeruginosa* biofilm formation is believed to be the main cause of persistent infections and tissue destruction. The PAO1/p_*lac*_-*yhjH* strain formed irregular, smaller cell aggregates at the infection sites at 2 dpi, with phagocytes penetrating the *P. aeruginosa* aggregates (Figure 3C). As a consequence of the phagocyte penetration, there were very few PAO1/p_*lac*_-*yhjH* cells remaining at 2 dpi compared with the corneas infected with PAO1 and Δ*wspF* (Figure 3A, B). A large number of phagocytes accumulated at the PAO1/p_*lac*_-*yhjH* infection sites at 7 dpi, with very few visible and viable PAO1/p_*lac*_-*yhjH* cells (Figure 3F). Determination of the bacterial load of *P. aeruginosa* strains at 2 dpi and 7 dpi correlated well with the imaging analysis and showed that c-di-GMP was required by *P. aeruginosa* for long-term colonization during corneal infection (Figure 3G).

**Figure 3.**
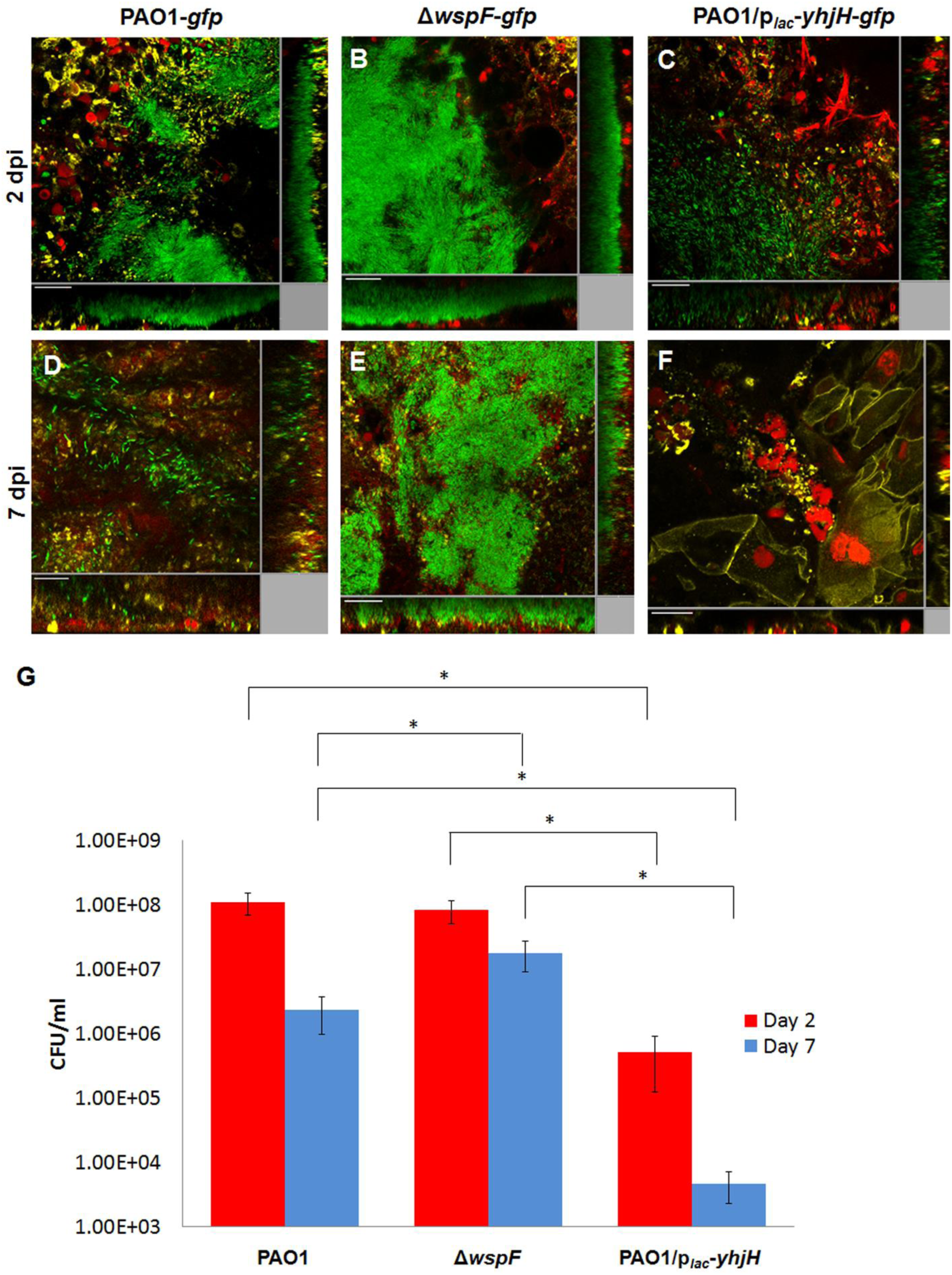
Distribution of *P. aeruginosa* cells and host cells among the *P. aeruginosa*-infected mouse corneas (A) and bacterial load determination reflected by colony forming units (CFU) (B) at 2 dpi and 7 dpi. Mouse corneas were infected with PAO1-gfp, Δ*wspF-gfp* and PAO1/p_lac_-*yhjH-gfp* strains. *P. aeruginosa* cells were tagged with *p_lac_-gfp.* The red fluorescence represents the staining of phagocyte lysosomes, including polymorphonuclear leukocytes, by LysoTracker® Red DND-99. The yellow fluorescence represents Alexa Fluor® 635 phalloidin, which stains F-actin of the eukaryotic cells. Representative confocal images are shown for each condition. Scale bars, 20 μm. CFUs of corneas infected with *P. aeruginosa* PAO1-gfp, Δ*wspF-gfp* and PAO1/p_lac_-*yhjH-gfp* at 2 dpi and 7 dpi. Mean values and s.d. from triplicate experiments are shown. *p ≤ 0.01, ANOVA test.

### Increased c-di-GMP content in P. aeruginosa modulates the host immune response

Biofilm formation has been proposed to shift bacterial infections from an acute phase to a chronic phase since biofilm cells can evade host immune attack and produce less acute virulence factors compared with planktonic cells [26,41,42]. To examine the host responses of mouse corneas toward *P. aeruginosa* infection, we examined the host-associated RNA reads using a dual RNA-Seq dataset to perform a comparative transcriptomic analysis of mouse corneas infected by the *P. aeruginosa* PAO1, Δ*wspF* and PAO1/p_*lac*_-*yhjH* mutants at 2 dpi and 7 dpi. Between 25,457,626 and 32,332,158 reads were mapped to the mouse genome (Table S1).

Projection of the host transcriptomes by PCA revealed that *P. aeruginosa* infections affected host gene expression in a time-dependent manner, during which 8 of 9 host transcriptomes grouped together at 2 dpi independently of the c-di-GMP content of the *P. aeruginosa* cells (Figure 4A). Interestingly, c-di-GMP was shown to affect host gene expression at 7 dpi, with a significant effect on PC1 genes observed in the high intracellular c-di-GMP strain (Figure 4A).

**Figure 4.**
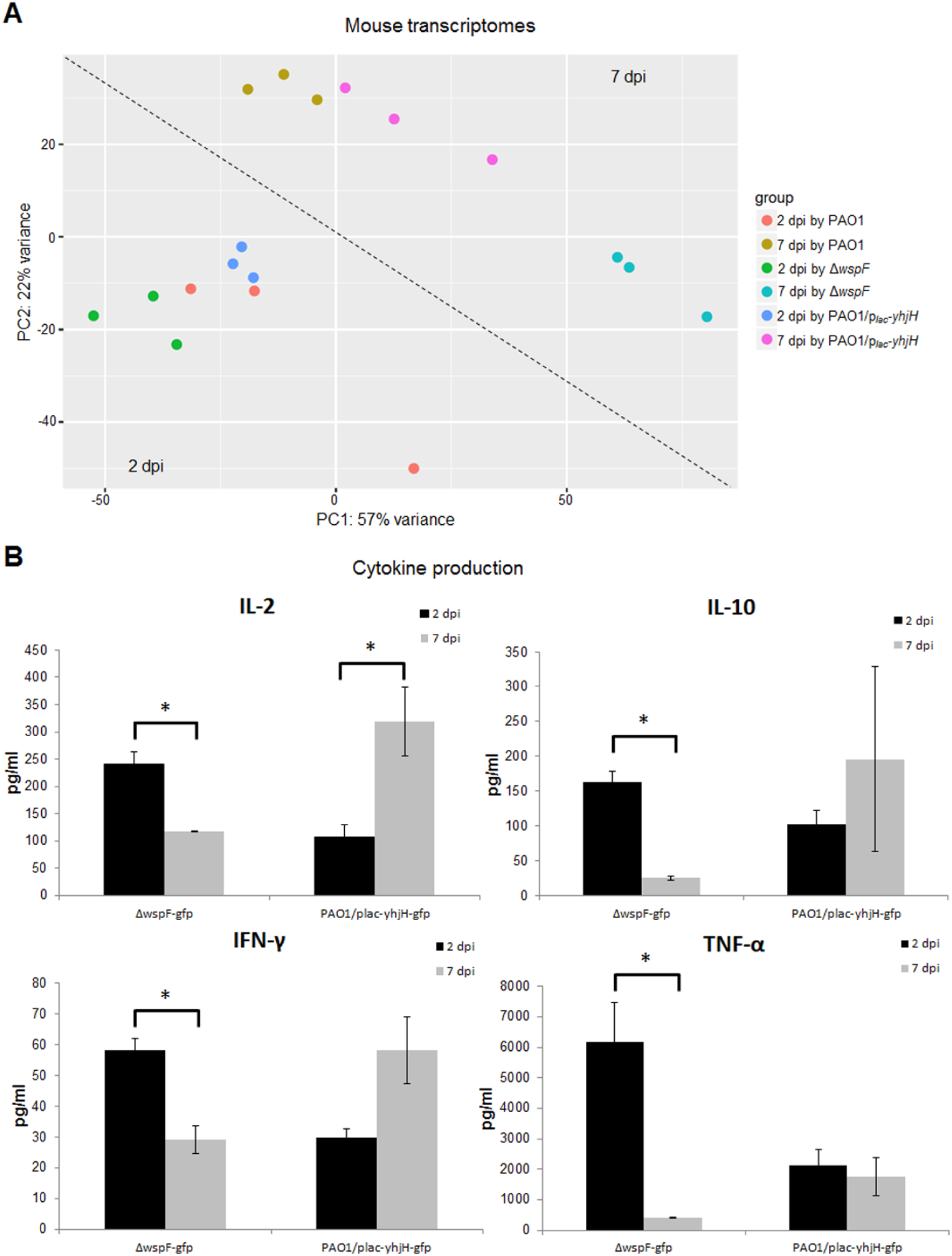
**A. PCA of transcriptomes of mouse corneal cells at 2 dpi and 7 dpi with *P. aeruginosa* PAO1, Δ*wspF* and *PAO1/p_lac_-yhjH.*** Raw RNA-Seq data were normalized using the DESeq package before PCA analysis. The dotted line was used to separate the transcriptomes of 2 dpi cells and *7* dpi cells by PCA analysis. **B. Production of cytokines by mouse corneal cells at 2 dpi and 7 dpi with *P. aeruginosa* Δ*wspF* and PAO1/p_*lac*_-*yhjH.*** Black bars represent 2 dpi and gray bars represent 7 dpi. Mean values and s.d. from triplicate experiments are shown. *p ≤ 0.05, **p ≤ 0.01, ANOVA test.

To further examine the role of c-di-GMP signaling in the modulation of host immune responses, we compared the host transcriptomes after infection by the two c-di-GMP ‘locked’ strains: Δ*wspF* and PAO1/p_*lac*_-*yhjH*. We excluded the host transcriptome of *P. aeruginosa* PAO1-infected corneas because of the potential fluctuation of c-di-GMP content of PAO1 during the infection (Figure 2I) due to the dynamic activities of different DGCs and PDEs.

Using a P-value cut-off of 0.05 and a fold-change cut-off of 5, 1385 host genes (Supplementary Data 5) were up-regulated and 507 host genes (Supplementary Data 6) were down-regulated in corneal cells at 7 dpi compared to corneal cells at 2 dpi caused by the Δ*wspF* mutant but not the PAO1/p_*lac*_-*yhjH* mutant. Thus, the expression of these host genes was specifically associated with increased c-di-GMP content in *P. aeruginosa*, and analysis of these genes might reveal important mechanisms underlying how the high intracellular c-di-GMP-containing clinical isolates (such as the small colony variant, SCV) evolved during chronic infections.

Several well-known genes involved in the host immune response toward bacterial infections were down-regulated, including *il10* (encoding interleukin 10) by 21.4-fold, *il23a* (encoding interleukin 23, alpha subunit) by 20.4-fold, *tnf* (encoding tumor necrosis factor) by 18.4-fold, and *Ifnlr1* (encoding interferon lambda receptor 1) by 9.8-fold, in corneal cells at 7 dpi compared with corneal cells at 2 dpi (Supplementary Table 8) by the Δ*wspF* mutant but not the PAO1/p_*lac*_-*yhjH* mutant. This result suggests that components of the Th1 and Th2-mediated immune responses were down-regulated during the late stage of corneal infection due to increased c-di-GMP content in *P. aeruginosa.*

To validate the transcriptomic analysis, C57BL6 mouse corneas were infected with the *P. aeruginosa ΔwspF* mutant and the PAO1/p_*lac*_-*yhjH* mutant. Infected mouse corneas were harvested at 2 dpi and 7 dpi to compare their cytokine levels. In accordance with the transcriptomic analysis, the cytokines IL-10, IFN-ƴ and transforming growth factor TNF-ɑ were all induced at 2 dpi in response to both *P. aeruginosa* strains (Figure 4B); these cytokines were reduced at 7 dpi in corneas infected with the Δ*wspF* mutant (Figure 4B), while they remained at the same levels or increased at 7 dpi in corneas infected with the PAO1/p_*lac*_-*yhjH* strain (Figure 4B). The reduced cytokine levels in mouse corneas infected with the *P. aeruginosa* Δ*wspF* mutant at 7 dpi compared with 2 dpi corresponded well with the very few phagocytes associated with the Δ*wspF* microcolonies at 7 dpi (Figure 3E). In contrast, the high cytokine levels in mouse corneas infected with the *P. aeruginosa* c-di-GMP-depleted PAO1/p_*lac*_-*yhjH* strain at 7 dpi compared with 2 dpi corresponded well with the large numbers of live phagocytes at the sites of infection at 7 dpi (Figure 3F). The cytokine levels have been previously reported to peak at early stages (1-3 d) of *P. aeruginosa* mouse corneal infection and then decline to baseline levels in BALB/c mice and remain at lower levels in C57BL/6 mice at the late stage of corneal infection (5-7 d) [43]. Our study showed that increased c-di-GMP content in *P. aeruginosa* was associated with the reduction of cytokine levels in the infected corneas of C57BL/6 mice, which suggests that the manipulation of intracellular c-di-GMP content in *P. aeruginosa* could be further investigated for improving current therapeutic strategies against biofilm-associated infections.

## Discussion

It has been proposed for decades that bacterial biofilm formation is responsible for many persistent and chronic infections. *In vitro* experiments have demonstrated that biofilms confer tolerance toward antimicrobials and phagocytosis by immune cells [41]. Numerous chemical biological strategies have also been developed to eradicate biofilms by targeting specific signaling pathways, such as quorum-sensing and c-di-GMP signaling, which are essential for *in vitro* biofilm formation [44]. However, the development of biofilm controlling strategies is largely limited by the lack of characterization of biofilms *in vivo.*

Using a mouse keratitis model, we have provided physiological evidence that *in vivo* biofilms formed by *P. aeruginosa* contribute to the establishment of ocular infections. Comparative transcriptomic analysis revealed that *P. aeruginosa* ocular infections induced Th17-regulated innate immune responses (Supplementary Data 5), which are known to contribute to the establishment of experimental *P. aeruginosa* infection in mouse corneas [45]. Importantly, we demonstrated that the bacterial secondary massager c-di-GMP plays a crucial role in the establishment of mouse corneal infections. A p_*cdrA*_-*gfp* reporter revealed the kinetics of the intracellular c-di-GMP content of *P. aeruginosa* during mouse ocular infections, in which the intracellular c-di-GMP level increased after 2 h of infection (Figure 2D) and declined at 7 dpi (Figure 2I). The induction of c-di-GMP signaling in *P. aeruginosa* during the early stage of ocular infections might be due to the down-regulation of several PDEs (Figure S1), which can lead to an accumulation of c-di-GMP. The induction of intracellular c-di-GMP content is crucial for the survival of *P. aeruginosa* during the early stage of mouse ocular infections. The PAO1/p_*lac*_-*yhjH* strain was cleared by the mouse immune system much more efficiently than the PAO1 wild-type and Δ*wspF* mutant strains (Figure 3). In contrast, lowering the intracellular c-di-GMP in PAO1/p_*BAD*_-*yhjH* and PAO1 at 1 dpi had no impact on the survival of these strains during infection (Figure 7). Similarly, a recent report has shown that c-di-GMP synthesis is also required for the establishment of *P. aeruginosa* biofilms *in vivo* during catheter-associated urinary tract infection [46].

We also observed a reduction of *P. aeruginosa* intracellular c-di-GMP content at 7 dpi, which might have resulted from the production of a large number of antimicrobial components (such as NO) during the course of infection. Unfortunately, due to the technical limitations of our RNA-Seq analysis, we were unable to obtain enough reads and identify potential PDEs responsible for the reduction of c-di-GMP at 7 dpi.

High c-di-GMP-containing *P. aeruginosa* variants (such as the Δ*wspF* mutant) are often isolated from chronic *P. aeruginosa* infections [33]. We provide in *vivo* evidence that a high c-di-GMP-containing *P. aeruginosa* variant is able to form large microcolonies during ocular infections (Figure 3). Phagocytic cells accumulated around but were unable to penetrate the *P. aeruginosa* biofilm microcolonies in the infected mouse corneas (Figure 3), in a similar manner to previously demonstrated *P. aeruginosa* microcolonies in infected lung tissue of cystic fibrosis patients [40]. Phagocytic cells that are recruited to infection sites are likely to cause severe damage during the host inflammatory response [43]. However, our study provides evidence that the *P. aeruginosa ΔwspF* mutant containing high levels of c-di-GMP also down-regulated Th1/Th2-regulated innate immune responses during the late stage of mouse corneal infection (Supplementary Data 6). C-di-GMP is known to control the production of Pel, Psl exopolysaccharides, CdrA adhesin, Cup fimbriae, and the pyoverdine iron siderophore and to decrease bacterial motility *in vitro* [19,47,48]. These biofilm-associated properties may impair host immune system clearance. For example, *P. aeruginosa* Psl polysaccharide production and reduced motility have been shown to be linked to a reduction in neutrophil opsonophagocytosis [49,50]. Interestingly, through comparative transcriptomic analysis of *in vivo* PAO1 at 2 dpi of the mouse corneas with planktonic cells, we noticed that only Pel polysaccharide but not Psl polysaccharide synthesis was induced *in vivo* (Supplementary Data 3). This result might partially explain why Psl polysaccharide is not produced by this widespread *P. aeruginosa* PA14 linage [51]. It remains unclear how Pel polysaccharide contributes to the tolerance of *P. aeruginosa* biofilms toward host immune clearance and antibiotic treatment. Further studies are needed to evaluate the Pel polysaccharide and other potential c-di-GMP effectors that contribute to the down-regulation of Th1/Th2-regulated innate immune responses.

## Acknowledgments

This research was supported by the National Research Foundation and the Ministry of Education of Singapore under its Research Centre of Excellence Programme, AcRF Tier 2 (MOE2014-T2-2-172 & MOE2016-T2-1-010) from the Ministry of Education, Singapore. S.L.C. was supported by the LKCMedicine Postdoctoral Fellowship.

## Contributions

J.K.H.Y. and T.T.A. designed the methods and experiments, carried out the laboratory experiments, analyzed the data and interpreted the results. G.S.K., F.C. and Y.L. co-designed the RNA sequencing experiments and performed the data analysis. D.D.M. and S.C.S. co-designed and performed the sequencing experiment and associated data collection. C.Z., J.Z. and Y.C. constructed the bacterial strains. S.A.R. and S.K. co-designed the biofilm-dispersing experiment. M.M.S. and S.L.C. co-designed and conducted the animal experiments. S.K., M.G., B.W., R.W.B. and T.T.N. discussed the analyses, interpretation and presentation. L.Y. wrote the paper. All authors have contributed to, seen and approved the submitted version of the manuscript.

## Competing interests

The authors declare that they have no competing financial interests.

